# UVB radiation suppresses Dicer expression through β-catenin

**DOI:** 10.1101/2023.12.22.572980

**Authors:** Zackie Aktary, Valérie Petit, Irina Berlin, Nisamanee Charenchaon, Evelyne Sage, Juliette Bertrand, Lionel Larue

**Affiliations:** Institut Curie, PSL Research University, INSERM U1021, Normal and Pathological Development of Melanocytes, Orsay, France; Université Paris-Saclay, Univ Paris-Saclay, CNRS UMR 3347, Orsay, France

**Keywords:** β-catenin, Dicer, UVB, transcription, melanocytes, melanoma

## Abstract

Ultraviolet (UV) rays prompt a natural response in epidermal cells, particularly within melanocytes. The changes in gene expression and related signaling pathways in melanocytes following exposure to UVR are still not entirely understood. Our findings reveal that UVB irradiation suppresses the expression of Dicer. This repression is intricately linked to the activation of the PI3K, RSK, and WNT/β-catenin signaling pathways and is directly associated with transcriptional repression by β-catenin. Notably, we have identified specific binding sites for the LEF/β-catenin complex in the Dicer promoter. Collectively, these results emphasize the significance of the UV-induced pathway involving LEF/β-catenin, which impacts Dicer expression. This pathway holds potential importance in governing melanocyte physiology.

## INTRODUCTION

Gene regulation is a central process in the life of cells, as well as in communication between cells and their (micro-)environment. Among the key players in this intricate network is Dicer, an enzyme that plays a pivotal role in governing the production of small RNA molecules crucial for modulating gene expression. It has been demonstrated that Dicer’s activity is subject to regulation by various factors, including reactive oxygen species (ROS), hypoxia, serum, interferons, and phorbol esters (Asada et al. 2008; van den Beucken et al. 2014; Wiesen and Tomasi 2009).

Dicer is a multifaceted enzyme belonging to the ribonuclease III (RNase III) family and acts as a conductor of gene expression by finely adjusting the levels of small RNA molecules, including microRNAs (miRNAs) and small interfering RNAs (siRNAs). These small RNA molecules, typically composed of 20-30 nucleotides, play a critical role in post-transcriptional gene regulation, dictating the fate of target mRNAs by either promoting their degradation or inhibiting their translation (Bartel 2018). Under conditions of oxidative stress, there is a surge in ROS levels, triggering a cascade of cellular responses. Among these responses is the downregulation of Dicer expression and activity, resulting in altered miRNA processing and subsequent changes in gene expression patterns (Chakraborty et al. 2017; Grishok et al. 2001; Kozomara and Griffiths-Jones 2014; Wiesen and Tomasi 2009). Phorbol esters, capable of mimicking the effects of diacylglycerol (DAG), a potent activator of protein kinase C (PKC), can induce the phosphorylation of Dicer, leading to enhanced miRNA processing and subsequent alterations in gene expression profiles (Shin et al. 2009). However, they can also repress the expression of Dicer (Wiesen and Tomasi 2009). Additionally, the presence of double-stranded RNA (dsRNA) and interferon-α (IFN-α) has been found to suppress Dicer production, while interferon-γ (IFN-γ) induces it. Serum withdrawal and hypoxia downregulate Dicer, increasing susceptibility to apoptosis (Asada et al. 2008; van den Beucken et al. 2014; Wiesen and Tomasi 2009).

In the context of skin biology, Dicer has been shown to play a pivotal role in essential aspects of hair follicle and epidermal morphogenesis and maintenance (Andl et al. 2006; Vishlaghi and Lisse 2020). In melanocytes, Dicer has been associated with apoptosis and disruption of proper melanocyte migration within the growing hair follicle, ultimately leading to the depletion of the pool of melanocyte stem cells (McSC) (Bertrand et al.; Levy et al. 2010). Epidermal cells, which are constantly exposed to the potentially harmful effects of ultraviolet radiation (UV) from sunlight, have evolved an array of defense mechanisms to counteract the detrimental impact of this radiation. These protective strategies encompass processes such as pigmentation, cell cycle inhibition, replication suppression, DNA repair, and apoptosis. Dysfunctions in these protective mechanisms can result in a range of pathophysiological conditions, including premature skin aging and manifestations of sun damage, such as wrinkles, leathery skin, liver spots, actinic keratosis, solar elastosis, loss of pigmentation, and the development of melanoma, a highly aggressive and increasingly prevalent form of skin cancer. UV exposure has been definitively linked to the emergence of skin conditions, including melanoma (Gorman et al. 2019), with epidemiological studies suggesting that a substantial proportion of melanoma cases, ranging from 65% to 90%, can be attributed to UV exposure (Glanz et al. 2007). Within the solar spectrum, UVB light (ranging from 290 to 320 nm) has been identified as a pivotal factor in the induction of melanoma (De Fabo et al. 2004; Gaffal et al. 2011; Mitchell and Fernandez 2011). Furthermore, Dicer’s involvement in the stress response and its regulation under various cellular and environmental conditions underscore the importance of understanding the context-specific modulation of Dicer expression. However, the precise mechanism governing the control of Dicer expression remains elusive. Given the critical role of melanocytes in shielding the skin against the harmful effects of UV radiation, this study aimed to investigate the regulation of Dicer gene expression in response to UV stress within the melanocyte lineage.

## RESULTS

### UVB represses Dicer at the transcriptional level

Given that UVB radiation primarily affects epidermal cells, we determined the effect of UVB radiation on Dicer expression in melanocytes. Dicer mRNA levels, as determined by RT-qPCR, were decreased when exposed to UVB radiation in melan-a mouse melanocyte cells in a dose-and time-dependent manner (Figure 1A-C). Further experiments were performed using 25 or 100 mJ/cm^2^ and the level of Dicer was determined 15 hours after irradiation. The level of Dicer mRNA was decreased across various cell lines after UVR, encompassing mouse melanocytes (melan-a, 9v), fibroblasts (NIH3T3) and keratinocytes (XB2); as well as human melanoma cells including 501Mel, MNT-1, and WM852 cells (Figure 1C). As expected, the level of Dicer protein was decreased across transformed and non-transformed cell lines (Figure 1D). These findings indicate that the reduction in Dicer expression following UVB exposure is not specific to the melanocyte lineage and is species independent. This decline in Dicer protein and mRNA levels after melanocyte exposure to UVB suggested that Dicer transcription is affected in response to UVB irradiation. To further explore this mechanism, we performed reporter assays in melan-a and MNT-1 cells with a luciferase reporter construct driven by the Dicer promoter. The results indicated that Dicer promoter activity was halved following UVB irradiation compared to mock-treated cells, providing evidence that the response of Dicer to UVB irradiation occurs, at least partially, at the transcriptional level (Figure 1E). These findings indicate that the regulation of Dicer transcription following UVB exposure is species independent.

**Figure 1.**
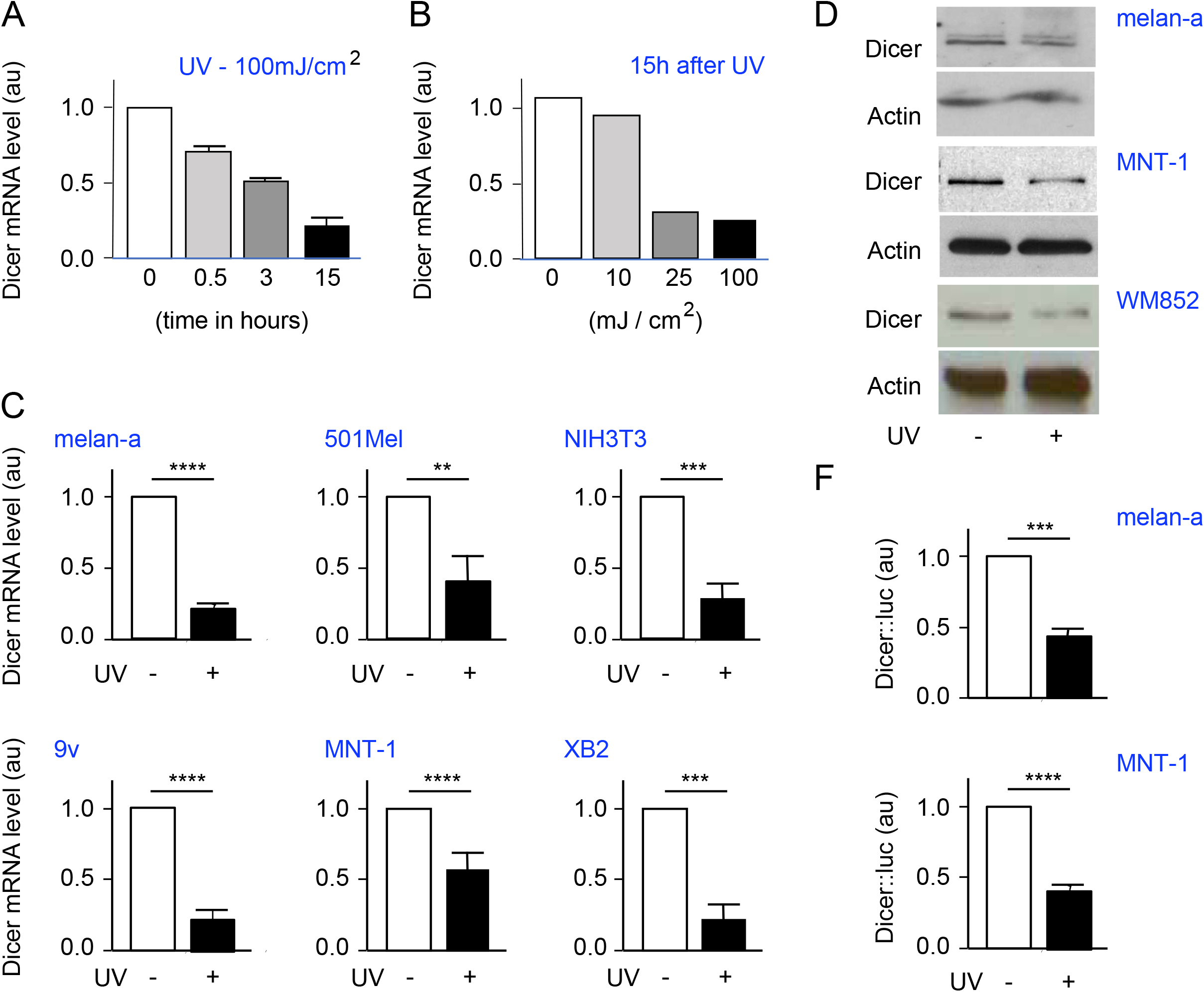
UVB-irradiation represses Dicer expression in various cell types. (A-C) Dicer transcript levels were assessed using RT-qPCR following UV irradiation or without UV exposure. (A)RNAs were extracted from melan-a cells 0-15 hours after UVB irradiation at 100 mJ/cm^2^. (B)RNAs were isolated from melan-a cells 15 hours after UVB irradiation ranging from 0 to 100 mJ/cm^2^. (C)RNAs were collected 15 hours after UVB irradiation at 100 mJ/cm^2^ (+) or without irradiation (-) from melan-a and 9v mouse melanocytes, and NIH3T3 mouse fibroblasts, and XB2 keratinocytes. Similar experiments were performed with 501Mel and MNT-1 human melanoma cells after UVB irradiation at 25 mJ/cm^2^. (D)Dicer protein levels were analyzed by Western blotting 15 hours after UVB irradiation: 100 mJ/cm^2^ for melan-a cells, and 25 mJ/cm^2^ for MNT-1 and WM852 cells. A representative Western blot is presented, with actin used as the loading control. (E)Dicer luciferase transcriptional activity was measured 15 hours after UVB irradiation (+) in melan-a (100 mJ/cm^2^) or in MNT-1 cells (25 mJ/cm^2^) or (-) no irradiation. Bars represent means ± SEM. Experiments were conducted independently at least three times, and statistical analysis was performed using an unpaired student t-test. Significance levels are denoted as follows: * p < 0.05; ** p < 0.01; *** p < 0.001; **** p < 0.0001.

### Dicer is regulated by UV through the GSK3β/β-catenin pathway

To determine which pathways regulate Dicer within the melanocyte lineage, we treated MNT-1 cells with pharmacological inhibitors targeting various signaling pathways (MAPK, PI3K, ATM, PKC, RSK) before assessing the level of endogenous Dicer mRNA. We assessed the effectiveness of each inhibitor by evaluating their impact on their known targets by Western blot analysis. MEK inhibitors (U0126), ATM inhibitors (KU55933), and PKC inhibitors (Enzastaurin) did not affect Dicer mRNA levels (Figure 2A-C). Conversely, PI3K (LY294002) and RSK inhibitors (BI-D1870) reduced Dicer mRNA levels to 0.6 and 0.5, respectively, compared to untreated controls (Figure 2D,E). Therefore, PI3K and RSK proteins play a role in regulating Dicer Mrna expression. Interestingly, both the PI3K and RSK pathways indirectly induce the inhibitory phosphorylation of GSK3β at Serine 9 (De Mesquita et al. 2001; Fang et al. 2000). Treating MNT-1 cells with a GSK3β inhibitor, BIO, reduced Dicer mRNA levels to 0.7 (Figure 2F), demonstrating that GSK3β also regulates Dicer mRNA levels.

**Figure 2.**
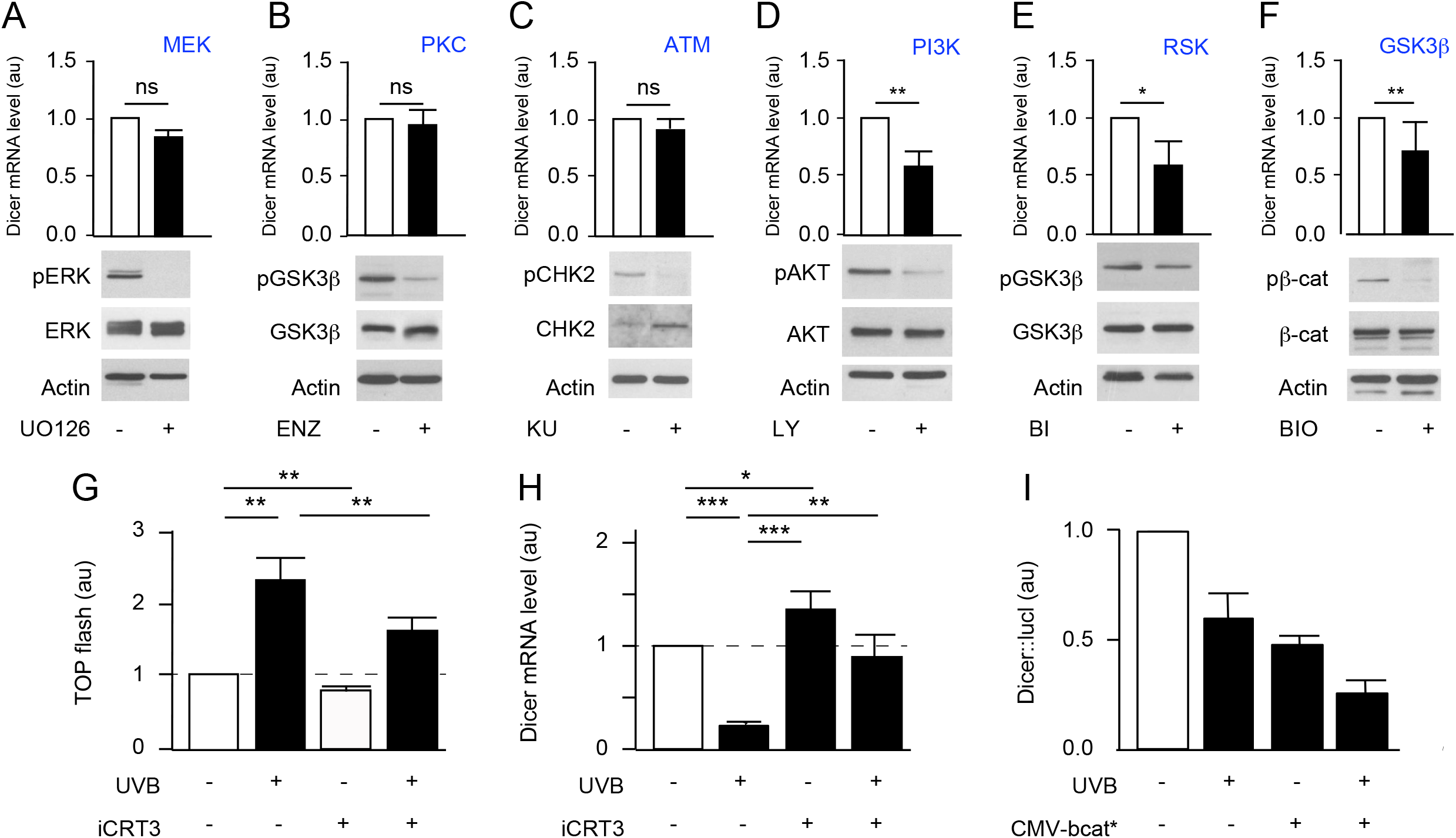
Regulation of Dicer via the PI3K, RSK, and GSK3β Pathways. Levels of endogenous Dicer mRNA in human melanoma MNT-1 cells after treatment or without treatment (DMSO) with inhibitors of MEK (A), PKC (B), ATM (C), PI3K (D), RSK (E), and GSK3β (F) are depicted. The Ct values obtained through quantitative PCR were normalized to the GAPDH housekeeping gene and further normalized to the untreated control using the ΔΔCt method. The inhibitors were used at the following concentrations and durations: (A) MEK inhibitor U0126, 10μM for 6 hours; (B) PKC inhibitor Enzastaurin, 10μM for 6 hours; (C) ATM inhibitor KU55933, 10μM for 24 hours; (D) PI3K inhibitor LY294002, 50μM for 6 hours; (E) RSK inhibitor BI-D1870, 20μM for 6 hours; (F) GSK3β inhibitor BIO, 5μM for 6 hours. The effectiveness of each inhibitor was validated by assessing the phosphorylation levels of downstream proteins: p(Thr202/Tyr204)-ERK (or p-ERK) for MEK, p(Ser9)-GSK3β (or p-GSK3β) for PKC and RSK, p(Thr68)-CHK2 (or p-CHK2) for ATM, p(Ser473)-AKT (or p-AKT) for PI3K, p(Ser33,Thr41,Ser45)-β-catenin (or p-βcat) for GSK3β. Actin was used as a loading control. (G) The efficacy of iCRT3 was validated by assessing TOP flash activity (Korinek et al., 1997). (H) Effect of UVB and/or iCRT3 on Dicer mRNA levels as assessed by RT-qPCR. (I) Effect of UVB and/or β-catenin expression on Dicer luciferase transcriptional activity. The mean and standard deviation are presented for at least three independent experiments (A-H). Statistical analysis was performed using a t-test: *** p-value< 10^-3^, ** p-value< 10^-2^, * p-value< 5 10^-2^, ns: not significant.

The best documented signaling function of GSK3β kinase is its phosphorylation of β-catenin at Serines 33, 37, and Threonine 41, leading to its subsequent inactivation (Peifer et al. 1994). Active β-catenin, in conjunction with transcription factors, particularly from the LEF/TCF family, regulates transcription. To determine whether GSK3β-mediated regulation of Dicer mRNA expression was modulated by β-catenin, MNT-1 cells were treated with an inhibitor of the transcriptional activity of the TCF-β-catenin complex, iCRT3, and validated by TOP flash analysis (Figure 2G) before analyzing the endogenous expression of Dicer mRNA. The iCRT3 inhibitor increased Dicer mRNA to 1.3 compared to the untreated control (Figure 2H). Furthermore, β-catenin expression and/or UVB reduced the level of Dicer transcription activity (Figure 2I). Hence, both GSK3β and β-catenin proteins participate in the regulation of Dicer mRNA expression. This hypothesis gained further support from the data set previously published (Bottomly et al. 2010), which indicated that β-catenin was capable of binding to the promoter region of Dicer.

### LEF/β-catenin binds to Dicer promoter and regulates its transcription

Because β-catenin is a recognized gene expression regulator, we investigated the Dicer promoter for potential TCF/LEF binding sites within the initial 300 base pairs upstream of the TSS (transcription start site) using the TRANSFAC database. ChIP experiments showed that both β-catenin and LEF-1 were associated with the Dicer promoter in SW620 cells (Figure 3A).

**Figure 3.**
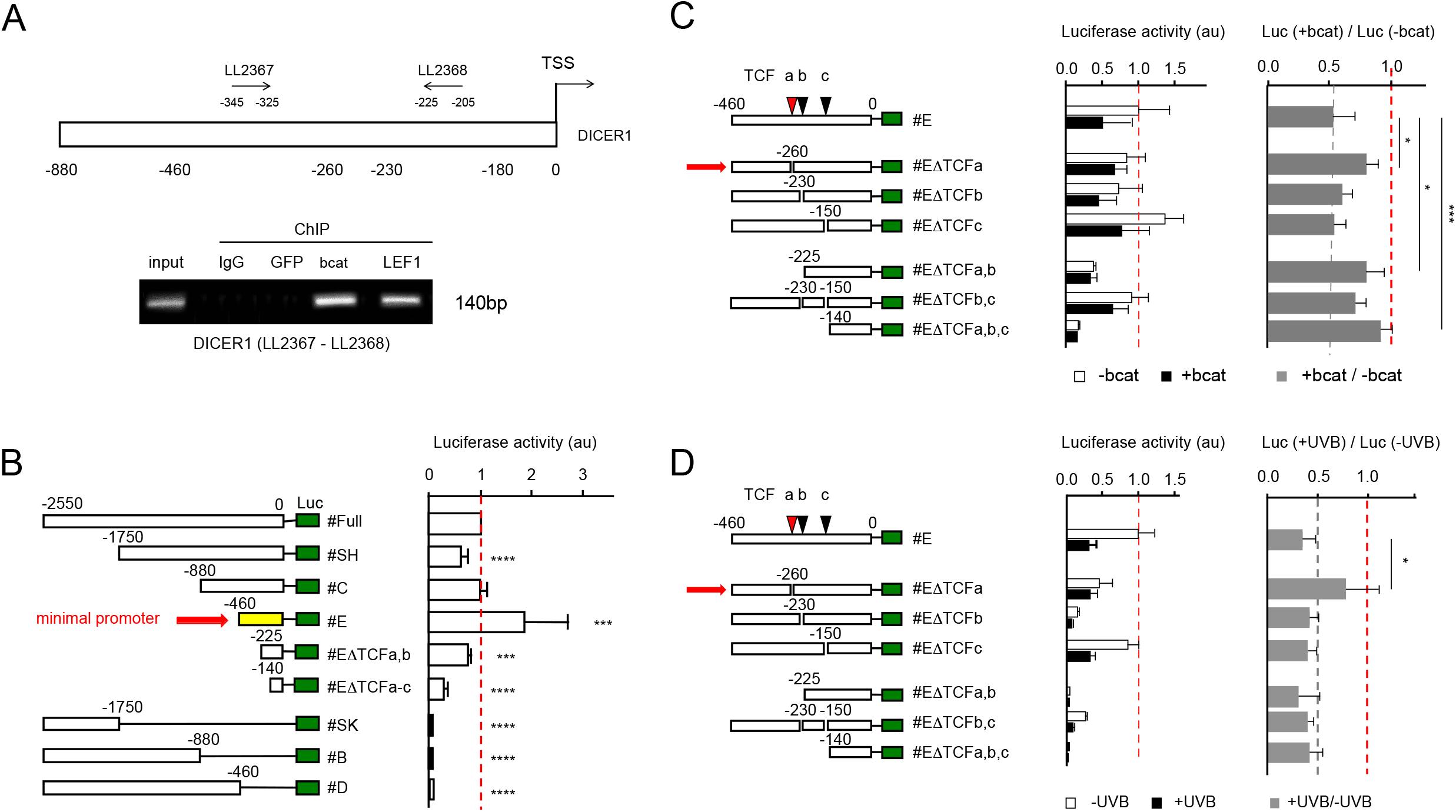
LEF/β-catenin directly regulates transcription of Dicer. (A)ChIP assays were conducted to assess the binding of β-catenin to the Dicer promoter in SW620 cells. The experiments were repeated three times (n=3). (B) A schematic representation illustrating various Dicer promoter constructs with either 5’ or 3’ regions removed. Luciferase activities (measured in arbitrary units, au) were assessed in melan-a cells to identify the minimal promoter region (indicated by the horizontal red arrow). (C, D) Schematics depicting various Dicer minimal promoter constructs with or without three potential LEF/β-catenin binding sites (represented as colored triangles). (C) Luciferase activities were measured in the presence or absence of exogenous β-catenin, revealing that constructs #E∆TCFa, #E∆TCFa,b, and #E∆TCFa,b,c exhibit reduced sensitivity to the presence of beta-catenin. (D) Luciferase activities were measured in the presence or absence of UVB irradiation, demonstrating that construct #E∆TCFa loses its sensitivity to UVB exposure. All experiments were independently conducted at least three times, and statistical analysis was performed using an unpaired Student’s t-test. Significance levels are indicated as follows: *p < 0.05, ***p < 0.001, and ****p < 0.0001. When not indicated, it signifies non significance.

To pinpoint the sequences/domains in the Dicer promoter necessary for regulating Dicer transcription, we generated truncated versions of the Dicer::luc promoter (Levy et al. 2010). We compared the luciferase activity of the full Dicer promoter with different truncated various in melan-a cells (Figure 3B). C-terminal deletions of the promoter (constructs #SK, #B, and #D) completely abolished luciferase activity, indicating the presence of essential elements near the TSS. Progressive N-terminal deletions of the promoter revealed that 460 base pairs upstream of the TSS still contained crucial elements for promoter activity. Thus, we designated construct #E as the minimal promoter (Figure 3B). Both the overexpression of β-catenin, and UVB irradiation resulted in decreased luciferase activity of the minimal promoter (Figure 3C-D).

*In silico* analysis of the minimal Dicer promoter sequence identified three putative binding sites for LEF/TCF, characterized as MAMAG (M=A/C) or CTKTK (K=G/T). To investigate the importance of these sites in regulating the Dicer minimal promoter, we generated deletions of the five nucleotides corresponding to these consensus sites, as described in the Materials and Methods section. Deletion of the TCFa binding site nearly abolished the β-catenin and UVB-mediated repression of luciferase reporter activity (Figure 3C,D, construct #EβTCFa). Individual deletions of the other two sites, TCFb and TCFc, or concomitant deletions of both, did not affect the luciferase activity response of the construct to β-catenin overexpression or UVB irradiation in a significant way. This analysis demonstrated that the repression of the Dicer promoter by β-catenin or UVB was dependent on the presence of TCF binding sites.

## DISCUSSION

The Dicer endonuclease is a key player in the regulation of miRNA biogenesis, and it plays a crucial role in various cellular processes. However, to date, only a limited number of upstream factors that modulate its activity have been identified (Asada et al. 2008; Levy et al. 2010; Martello et al. 2010; Su et al. 2010; Wiesen and Tomasi 2009). In this study, we have identified β-catenin as a regulator of Dicer expression in response to UV stimulation in melanocytes.

We previously demonstrated that UVB resulted in the nuclear accumulation of β-catenin and its subsequent transcriptional activity in mouse melanocytes (Bertrand et al. 2017). Our results here have demonstrated that this phenomenon is also observed in human melanoma cells. More significantly, we have shown that the Dicer gene is a transcriptional target of β-catenin/LEF/TCF and that its expression is downregulated, in a β-catenin-dependent manner, following UVB irradiation in the melanocyte lineage. While our results are the first demonstration of β-catenin regulating Dicer in melanocytes, the relationship between β-catenin and Dicer was shown in the HEYA8 ovarian cancer cell line, after conducting a classical functional analysis by both overexpressing and suppressing the level of β-catenin (To et al. 2017). However, the precise mechanism through which β-catenin represses Dicer remained unexplored at the time (To et al. 2017). Our results also suggest that UVB-mediated Dicer repression is not cell-type specific, since we observed similar results in melanocyte, keratinocyte and fibroblast cell lines.

Various factors, including reactive oxygen species (ROS), hypoxia, serum, interferons, and phorbol esters, regulate Dicer’s activity (Asada et al. 2008; van den Beucken et al. 2014; Wiesen and Tomasi 2009). Additionally, UV radiation, potentially UVC, has been shown to suppress Dicer mRNA expression in specific cells, like mouse preadipocytes (Mori et al. 2012). However, the underlying mechanism behind this suppression remained unreported. While the regulation of Dicer transcription has not been deeply studied, different transcription factors are thought to play a role in different cellular contexts. A previous report showed that in melanoma cells, SOX4 promotes Dicer expression by binding to the Dicer gene promoter (Jafarnejad et al. 2013). Loss of SOX4-mediated Dicer expression resulted in increased melanoma cell invasion (Jafarnejad et al. 2013). We also have observed increased melanocyte cell migration following decreased Dicer expression (Bertrand et al.). Interestingly, previous studies have also suggested that SOX4 competes with β-catenin for binding to TCF/LEF in colon carcinoma cells (Sinner et al. 2007), thus acting as a negative regulator of β-catenin transcriptional activity. In this respect, β-catenin and SOX4 may be involved in differentially regulating Dicer expression, at least in the melanocyte lineage.

In melanocytes, the bHLH leucine zipper transcription factor M-Mitf, which is specific to this lineage, has been identified as a positive regulator of Dicer transcription (Levy et al. 2010). Our data indicates that both Mitf and β-catenin, in association with Lef1, exhibit antagonistic roles in the transcriptional regulation of Dicer. As β-catenin positively regulates the transcription of Mitf, and Mitf directly interacts with Lef1 and/or β-catenin (Luciani et al. 2011; Saito et al. 2003; Schepsky et al. 2006; Takeda et al. 2000), it underscores the critical importance of regulating the quantities and activities of these two proteins to modulate Dicer levels in the melanocyte lineage. This intricate interplay between β-catenin and Mitf has been previously observed in other pathways. For example, β-catenin is known to positively regulate the transcription of Pou3f2 (Brn2), and Brn2, in turn, may regulate Mitf either positively or negatively, depending on the phosphorylation status of Brn2 and the implication of Pax3 (Berlin et al. 2012; Smith et al. 2019). Furthermore, a negative feedback loop operates from Mitf to Brn2 via mir-211 (Boyle et al. 2011).

Dicer expression is also regulated post-transcriptionally, as a number of alternative splicing variants have been identified for the Dicer transcript, with some encoding for shorter Dicer isoforms and others not encoding any protein (Grelier et al. 2009; Hinkal et al. 2011; Kurzynska-Kokorniak et al. 2015). A link may exist between β-catenin and alternative splicing. Indeed, it has been shown in colorectal cancer cell lines that β-catenin regulates the expression of the splicing factor SRp20 (Gonçalves et al. 2008). For now, we have not evaluated the potential role of SRp20 nor β-catenin in the alternative splice regulation of Dicer in melanocytes and melanoma, or whether UVB has an effect on the alternative splicing of Dicer transcripts.

Dicer is a crucial protein that is involved in a number of fundamental processes, including embryonic development and cellular stress responses, through the regulation of gene expression and modulation of DNA repair, respectively. As such, understanding the mechanisms regulating its expression is of vital importance. By identifying and characterizing β-catenin as a transcriptional regulator of Dicer following UVB radiation, our work brings to light the importance and intricacy of the network consisting of β-catenin, Dicer, Mitf, UVB, cell migration, pigmentation and DNA repair. Moreover, our results underscore the fundamental roles played by UVB, β-catenin, and Dicer in establishing, renewing, and transforming the melanocyte lineage.

## EXPERIMENTAL PROCEDURES

### Cell Culture and Transfection Procedures

The melan-a mouse melanocyte cell line was generously provided by Dr. D. Bennett (Bennett et al. 1987), and 9v mouse melanocyte cell line was previously generated (Delmas et al. 2007). These cells were cultured in F12 medium supplemented with 10% fetal calf serum, and 200nM TPA (tetradecanoyl phorbol acetate, Sigma). MNT-1, WM852 and 501mel human melanoma cells were grown in RPMI-1640 medium supplemented with 10% fetal calf serum (Moore et al. 2004). Murine NIH3T3 fibroblasts and XB2 keratinocytes were maintained in DMEM medium (GIBCO) supplemented with 10% fetal calf serum. SW620 human colon carcinoma cells were grown in Leibovitz’s L-15 media supplemented with 10% fetal calf serum and 2mM L-glutamine.

Antibiotics (100 U/mL penicillin and 100 μg/mL streptomycin) were added to all culture media, and cells were incubated at 37°C in a humidified atmosphere with 5% carbon dioxide.

For transient transfection, cells at 70% confluence were transfected using Lipofectamine 2000 (Invitrogen) following the manufacturer’s instructions. In luciferase assays, cells seeded in 12-well plates were co-transfected with either 1 μg of total plasmid DNA or 100nM siRNA, along with the TK::Renilla luciferase construct used as a control. The mass of DNA or siRNA was adjusted to be equalized with pBluescript. Firefly and Renilla luciferase activities were assessed 48 hours post-transfection, unless otherwise specified, using the Dual Luciferase Reporter Assay kit from Promega. Firefly luciferase activity was normalized against Renilla luciferase activity.

### UV Irradiation and Sample Preparation

UV irradiation was conducted using a VL-330 mid-range lamp with a continuous spectrum ranging from 250 to 400 nm, with a peak emission at 313 nm. The dose rate of our UV system underwent calibration by the French National Metrology Institute (LNE France -Laboratoire National de Metrologie et d’Essais). Based on the spectral energy distribution of the UV source, measured at a distance of 23 cm from the lamps, it was determined that 70% of the total radiation fell within the UVB range, 29.9% within the UVA range, and a minimal component (0.1%) resided at the upper end of the UVC range. The spectral output of the utilized source can be found in Supplementary Figure S1. To achieve a dose of 100 mJ/cm^2^ UVB, an exposure duration of 25 seconds was necessary. To ensure consistent UVB radiation levels, we regularly monitored UVB intensity using an IL1700 radiometer before each exposure, occurring every other week.

For UV irradiation of cell cultures, we first removed the medium from cultures when they reached 80% confluence. Subsequently, we washed the cells twice with PBS and replaced the medium with fresh PBS. During UV irradiation, the plastic dish’s lid was removed. Following UV exposure, we substituted the PBS with fresh culture medium, and the cells were then incubated for varying durations, depending on the specific experiment’s requirements. Control cultures underwent a similar treatment process but omitted the UV exposure step.

### RNA Extraction and RT-qPCR Analysis

To extract total RNA from cultured cell lines, we utilized the RNeasy mini kit (Qiagen), following the manufacturer’s guidelines. The recovered RNA was quantified using a Nanovue device (General Electric).

Reverse transcription reactions were conducted using 1 μg of total RNA and M-MLV reverse transcriptase (Invitrogen), following the manufacturer’s protocols. The resulting cDNA was then subjected to semi-quantitative real-time PCR analysis of gene expression. This reaction was performed in a QuantStudio™ Real-time PCR System. We estimated the RNA quantities corresponding to Dicer and Gapdh using the appropriate oligonucleotides and SYBR Green Supermix (Bio-Rad) (Table S1A). The thermal profile consisted of an initial denaturation step at 95°C for 90 seconds, followed by 40 cycles of denaturation at 95°C for 30 seconds and annealing/extension at 60°C for 1 minute. The obtained results were normalized to the expression of the Gapdh gene, which served as a housekeeping gene. The relative amount of the target transcript in comparison to the reference gene (GAPDH) transcript was determined using the ΔΔCt method. Each experiment was repeated a minimum of three times, and the results from triplicate experiments are presented for each time point.

### Western blotting procedure

To initiate Western blotting, cells were lysed in ice-cold RIPA buffer enriched with a complete protease inhibitor cocktail and PhoStop phosphatase inhibitor cocktail (Roche, France). The lysates were then cleared of cell debris by centrifugation at 14,000 rpm for 15 minutes. The resulting supernatants were collected, and their protein concentrations were determined using a BCA assay. Subsequently, 50-100 μg of total proteins were separated on a denaturing 10% acrylamide SDS-PAGE gel through electrophoresis.

Following electrophoretic separation, proteins were transferred onto a nitrocellulose membrane, which was further treated with Ponceau Red solution to assess the quality of the transfer. To block nonspecific binding, the membrane was incubated for 1 hour in a solution containing 5% nonfat milk powder in Tris-buffered saline supplemented with 0.01% Tween-20 (TBST). The membrane was then subjected to an overnight incubation with primary antibodies, dissolved in a 5% nonfat milk powder TBST solution. Afterward, the membrane underwent three rounds of washing with TBST.

Subsequently, the membrane was incubated with the corresponding horseradish peroxidase (HRP)-conjugated secondary antibodies at a 1:10,000 dilution. Following another set of three TBST washes, antibody binding was detected using enhanced chemiluminescence (ECL; Thermo, France). The list of primary antibodies used is provided in Table S1B, including Dicer, Actin, p204-ERK, ERK, p9GSK3b, GSK3b, p68CHK2, CHK2, p473AKT, AKT, p33,37,41bcat, and bcat. All experiments were independently conducted at least three times.

### Chromatin Immunoprecipitation (ChIP)

ChIP experiments were performed as previously described (Aktary et al., 2013). The composition of all buffers is provided in Table S1C. Confluent 150 mm cultures were trypsinized and 2x10^7^ cells pelleted by centrifugation at 3,500 rpm for 10 minutes. The cell pellets were then resuspended in growth media to which formaldehyde (Fisher) was added to a final concentration of 1% and incubated at room temperature for 10 minutes. To stop fixation, glycine was added to a final concentration of 125 mM. The cell suspension was then centrifuged at 3,500 rpm at 4°C for 10 minutes. The resulting cell pellets were then washed twice with PBS containing 1 μg/ml aprotinin and leupeptin and 1 mM PMSF, after which they were resuspended in cell lysis buffer and incubated on ice for 15 minutes. NP-40 was then added (final concentration of 0.6%) after which the samples were vortexed for 10 seconds at high speed and subsequently centrifuged at 13,000 rpm for 30 seconds. The resulting pellets were then resuspended in sonication buffer and left on ice for 10 minutes. The samples were then sonicated (Branson Sonifier 450) for 1 minute at 20% output for a total of four times.

The sonicated chromatin samples were then diluted ten-fold in chromatin dilution buffer after which 50 μl was removed (Input). Fourty μl Protein A/G Agarose beads (Calbiochem) were added and the samples were pre-cleaned on a rocker-rotator at 4°C for 2 hours. Following incubation, the samples were centrifuged briefly, and the resulting supernatant (pre-cleaned chromatin) was split into equal aliquots and processed for immunoprecipitation. Each aliquot was incubated with 5 μg antibodies and 40 μl pre-cleaned (by overnight incubation with 4 μg Salmon Sperm DNA and BSA) Protein A/G Agarose beads overnight at 4°C on a rocker-rotator.

Following immunoprecipitation, the samples were centrifuged for 10 minutes at 2,000 rpm at 4°C, after which the resulting supernatants were removed. The beads were then subjected to six 5-minute washes in each of four wash buffers. Following the washes, the protein-DNA complexes were eluted off the beads by incubation in elution buffer for 15 minutes at room temperature on a rocker-rotator.

Following elution, 1 μg RNase and NaCl (final concentration 300 mM) were added to the samples, which were then incubated at 65°C for 4 hours. Next, Tris pH 6.8, EDTA (final concentrations of 40 mM and 10 mM, respectively) and 4 μg proteinase K were added to the samples, which were incubated at 45°C for 2 hours. The samples were then purified using a PCR Purification Kit (QIAGEN, Valencia, CA) and processed for PCR.

### Plasmid constructs

The wild-type Dicer::luciferase promoter was kindly provided by Dr. D. Fisher and has been described elsewhere (Levy et al. 2010). The constructs #B, #C, #D, #E, #SH, #SK, #EΔTCFa,b,c of Dicer promoter were generated by digesting the full Dicer::luciferase promoter with the following restriction enzymes: *KpnI* and *NcoI* (#B), *NcoI* and *HindIII* (#C), *KpnI* and *EcoRI* (#D), *EcoRI* and *HindIII* (#E), *SmaI* and *HindIII* (#SH), *SmaI* and *KpnI* (#SK), *SacII* and *HindIII* (#EΔTCFa,b,c). The resulting fragments were eluted on agarose gel and inserted into the pGL3 luciferase vector digested by *SmaI* and *KpnI* or *SmaI* and *HindIII*.

The constructs #EΔTCFa, #EΔTCFb, #EΔTCFc, #EΔTCFa,b of Dicer promoter were generated by PCR-mediated deletion mutagenesis by overlap extension PCR in 2 steps to remove the putative binding site sequences: TCFa = CTTTC, TCFb = CACAG, TFCc = CTGTG. The oligonucleotides list is found in Table S1A.

## ACKNOWLEDGEMENTS

We thank David E Fisher for his generous gift of the Dicer::luciferase construct. Work in the laboratories is supported by grants from the CNRS, the INSERM, FRM, and INCa grant. LL is ‘équipe labellisée’ from FRM EQU202103012599. IB was supported by Institut Curie and LNCC fellowships. We are grateful to Morgane Verbrugghe for her technical assistance.

