## Supplementary material for "UVB radiation suppresses Dicer expression through β-catenin": Table S1A

**Table S1A. Oligonucleotides list**

| *RT-qPCR quantification* | | Primer forward | | Primer reverse | | Product size (bp) | Concentration |
|---|---|---|---|---|---|---|---|
| Species | Gene | Name | Sequence (5' - 3') | Name | Sequence (5' - 3') | | |
| Mouse | Dicer | LL1795 | AGC ATG GCA GGC CTG CAG | LL1786 | TGA TGG GCC AGC TCT TTG G | 224 | 300nM |
| Human | Dicer | LL1887 | CAA ATT TAG CCC GGC TGA GAG | LL1888 | GGC AAT CCT GTA ACT TCG ACC A | 106 | |
| Mouse | Gapdh | LL778 | ACC CAG AAG ACT GTG GAT GG | LL779 | CAC ATT GGG GGT AGG AAC AC | 171 | 300nM |
| Human | TBP | LL521 | CAC GAA CCA CGG CAC TGA TT | LL522 | TTT CTT GCT GCC AGT CTG GAC | 89 | 300nM |

| *ChIP* | | Primer forward | | Primer reverse | | Product size (bp) |
|---|---|---|---|---|---|---|
| | | Name | Sequence (5' - 3') | Name | Sequence (5' - 3') | |
| | | LL2367 | GCA AAC GCC TCT CCG GCC TC | LL2368 | GGC ATG AGA GCG AGC CTG TG | 120 |

| *Plasmids constructs* | |
|---|---|
| Name | (5' - 3') |
| LL408 | GTT CCA TCT TCC AGC GGA TA |
| LL2153 | AGA GCG AGC ATT GGA CAG GGC C |
| LL2154 | CTG TCC AAT GCT CGC TCT CAT GC |
| LL2158 | GGA ATG CCA AGC TTT CGT CC |
| LL2176 | AGC TAA GCT CTC CGG GAA AC |
| LL2210 | GGA TTC TCC AAG CGG CGC CGT TGC CGC |
| LL2211 | GCG GCA ACG GCG CCG CTT GGA GAA TCC |
| LL2261 | GCT CTC ATG CCC CGG GCG CCA CGG GG |
| LL2312 | GCC GGG ATT AAC ACC GCC AGG CCC TG |
| LL2313 | CAG GGC CTG GCG GTG TTA ATC CCG GC |
