## Supplementary material for "UVB radiation suppresses Dicer expression through β-catenin": Table S1B

**Table S1B. Antibodies list**

| Primary Antibodies | source | Identifier |
|---|---|---|
| Dicer | Abcam | ab13502 |
| Actin | Sigma | A5441 |
| ERK | CST | 9102 |
| pERK | Santa-Cruz | sc-7383 |
| GSK3b | Santa-Cruz | sc-9166 |
| pGSK3b | CST | 9336 |
| CHK2 | Santa-Cruz | sc-9064 |
| pCHK2 | CST | 2661 |
| AKT | CST | 4685 |
| pAKT | CST | 9271 |
| b-catenin | Abcam | ab6302 |
| pb-catenin | CST | 9561 |

| Secondary Antibodies | source | Identifier |
|---|---|---|
| anti-mouse HRP | Jackson Laboratory | 115-005-003 |
| anti-rabbit HRP | Jackson Laboratory | 111-035-003 |

| ChIP Antibodies | source | Identifier |
|---|---|---|
| b-catenin | CST | 9581 |
| LEF1 | Santa Cruz | sc-8592 |
| GFP | Institut Curie | |
| Normal Rabbit IgG | CST | 2729 |
