## Supplementary material for "UVB radiation suppresses Dicer expression through β-catenin": Table S1C

### Table S1C. ChIP buffer composition

| | Component | concentration |
|---|---|---|
| **Cell lyses buffer** | EDTA | 10 mM |
| | Tris HCl pH8.0 | 50 mM |
| | SDS | 1% |

| | Component | concentration |
|---|---|---|
| **ChIP dilution buffer** | SDS | 0,01% |
| | Triton X-100 | 1,10% |
| | EDTA pH8.0 | 1.2 mM |
| | Tris HCl pH8.0 | 16.7 mM |
| | NaCl | 167 mM |

| | Component | concentration |
|---|---|---|
| **Low salt wash buffer** | SDS | 0,10% |
| | Triton X-100 | 1% |
| | EDTA | 2 mM |
| | Tris pH 8.0 | 20 mM |
| | NaCl | 150mM |

| | Component | concentration |
|---|---|---|
| **High salt wash buffer** | SDS | 0,10% |
| | Triton X-100 | 1% |
| | EDTA | 2 mM |
| | Tris pH 8.0 | 20 mM |
| | NaCl | 500 mM |

| | Component | concentration |
|---|---|---|
| **LiCl wash buffer** | LiCl | 0.25 M |
| | NP40 | 1% |
| | Na deoxycholate | 1% |
| | EDTA | 1 mM |
| | Tris pH 8.0 | 10 mM |

| | Component | concentration |
|---|---|---|
| **Final wash buffer** | EDTA | 1 mM |
| | Tris pH 8.0 | 10 mM |

| | Component | concentration |
|---|---|---|
| **Elution Buffer** | SDS | 1% |
| | NaHCO3 | 0.1 M |
