## Supplementary figures and images for "UVB radiation suppresses Dicer expression through β-catenin"

### Figure S1

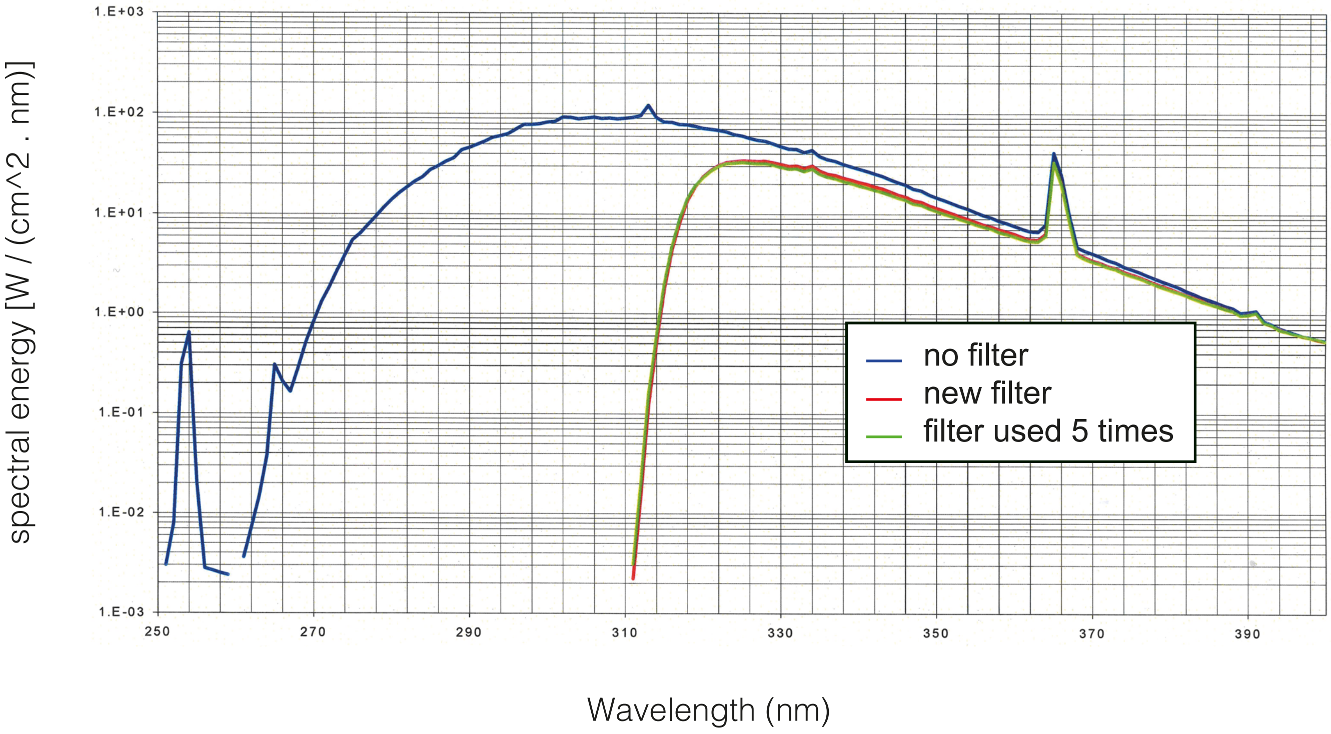
